# Spatiotemporal profiling of T cell repertoires in Lynch syndrome reveals convergent immune responses through neoplastic progression

**DOI:** 10.64898/2026.10.06.756982

**Authors:** Penelope Edwards, Ottilie Swinyard, Tahel Ronel, Yanrong Jiang, Gayathri Nageswaran, Chirine Sakr, Suhana Yasmin, Heather E Grant, Eleanor Smith, Konstantin Bräutigam, Sindhu Varughese, Pooja Datt, Adriana Martinez, Jennifer Fisher, Benny Chain, Kai-Keen Shiu, Marnix Jansen, Ann-Marie Baker, Kevin J Monahan, Trevor A Graham

## Abstract

Surveillance of individuals with Lynch syndrome (LS), who are carriers of heterozygous inactivating mismatch repair (MMR) gene mutations that predispose to colorectal cancer, enables the study of tumour-immune coevolution during tumour initiation. Somatic loss of MMR activity in the LS colon leads to the accrual of cancer driver mutations, and also of neoantigens likely to generate an adaptive immune response. However, the spatial and temporal immune dynamics in colon and peripheral blood are uncharacterised. Here we show that the neoplastic evolution of mismatch repair deficient (MMRd) clones leads to large-scale and sustained reorganisation of the T-cell repertoire in colon and peripheral blood. We performed dense spatiotemporal profiling of T cell receptor (TCR) repertoires from 109 colonic biopsies (11 tumour samples, 98 normal bowel) from seven individuals, with eight matched blood repertoires. We additionally analysed 13 colorectal polyp repertoires from eight participants, 286 cross sectional blood repertoires from 242 participants (198 LS and 44 non-LS) and 52 longitudinal peripheral blood samples from 26 individuals with LS. We detected tumour-associated T cell clones, some of which were widely present in normal bowel tissue and also detectable in peripheral blood. Sequence similarity analysis revealed conserved TCR architectures across tissue compartments and recurrent clustering patterns between patients, that were not present in non-Lynch controls, consistent with convergent MMRd-associated antigen-driven immune selection. We identified systemic signatures associated with local tumour immune activity and revealed evidence of structured repertoire remodelling before overt malignancy. Longitudinal profiling revealed frequent turnover of low-abundance clonotypes, whereas highly abundant clonotypes persisted for several years. Collectively, our data defines the local and circulating TCR landscape associated with colonic neoplasia in LS, and provides a framework for LS blood-based immune monitoring, early interception and biomarker development.

## Introduction

Lynch syndrome (LS) is a common (1 in 400 people) hereditary cancer predisposition syndrome where carriers have a heterozygous germline mismatch repair (MMR) gene mutation^1^. Somatic loss of the second wild-type allele leads to deficient MMR (dMMR) and the subsequent pathogenesis of cancers, typically in colon and endometrium. To mitigate cancer risk, LS patients are offered regular endoscopic bowel surveillance where colonic biopsies are collected. Surveillance therefore makes LS a unique human model in which the earliest interactions between emerging neoplastic clones and the adaptive immune system can be studied *in vivo* over space and time. Despite colonoscopic surveillance, colorectal cancers continue to arise in LS, including “interval cancers” which develop shortly after apparently normal examinations^2,3^. This suggests that cancer evolution in LS is an active process, where clonal dynamics can be relatively rapid.

Due to dMMR, cancers arising in LS carry a high burden of single nucleotide variants (SNVs) and short insertion-deletion (indel) mutations. A subset of these mutations leads to the production of neoantigenic peptides that are presented on the cell’s surface via the human leukocyte antigen (HLA), and consequently MMRd tumours are typically highly immunogenic. Further, patients often share recurrent frameshift neoantigens^4,5^.

Cancer evolution in LS therefore likely reflects a dynamic interplay between tumour emergence (due to the accrual of driver mutations), immune-mediated elimination (acting on the high burden of neoantigens) and the subsequent evolution of immune escape that enables cancer cells to grow despite carrying a high neoantigen burden. Immune architecture therefore is expected to attenuate neoplastic change but has not been studied directly.

The T-cell receptor (TCR) repertoire provides a functional record of antigen-driven immune responses^6^. Upon recognition of a cognate antigen, antigen-specific T cells undergo clonal expansion, generating measurable changes in repertoire organisation, including expansion of reactive clonotypes and the shifting of the repertoire towards T cells with related TCR clonotypes. Consequently, TCR repertoire analysis has emerged as a powerful, quantitative, approach for interrogating immune responses across infection, autoimmunity and cancer^7,8^. In tumour immunology, expanded and convergent TCR architectures are thought to reflect coordinated immune responses against shared antigenic stimuli and may provide insight into the spatial organisation of anti-tumour immunity.

In LS-associated neoplasia, the high neoantigen burden leads to marked T-cell infiltration and reactivity against frameshift peptide neoantigens^9^. Tumour-reactive T cells have been identified both within tumour tissue and in peripheral blood, suggesting that antigen-specific immune responses may arise early during neoplastic evolution. Recent studies have further demonstrated that circulating TCR signatures associated with dMMR neoplasia can be detected in cancer-free LS patients^10^, supporting the concept that systemic immune repertoires reflect ongoing immune surveillance activity.

The spatial organisation and temporal dynamics of these immune repertoires across tumour, normal bowel, polyp and circulating blood compartments in LS remain poorly understood. We hypothesised that the LS bowel harbours spatially structured convergent antigen-driven T-cell repertories that extend beyond tumour tissue and are detectable within the peripheral circulation, as a somatically-evolved response against recurrent neoantigenic stimuli arising during LS-associated carcinogenesis. We performed dense spatiotemporal profiling of TCR repertoires across tumour, polyp, histologically normal bowel and both longitudinal and single timepoint peripheral blood samples from individuals with and without LS and characterised the spatial organisation and systemic conservation of tumour-associated immune repertoires across the LS bowel ecosystem.

## Results

### Spatiotemporal profiling of LS patient colonic and blood samples

We analysed six colonic surgical specimens and one small bowel surgical specimen, all collected and processed at St Mark’s Hospital on the day of surgery. We performed dense spatiotemporal profiling of both normal and cancer colonic tissue (number of tissue samples: LS130 n=23, LS236 n=24, LS295 n=24, LS299 (small bowel) n=11, LS331 n=9, LS332 n=11, LS383 n=12), with paired peripheral blood samples where available (Figure 1A). We also collected 13 FFPE polyp samples from 8 patients, with 6 paired peripheral blood samples. The total tissue repertoires passing quality control was n=124 (cancer n=11; normal n=97; polyp n=13) (Figure 1C).

**Figure 1.**
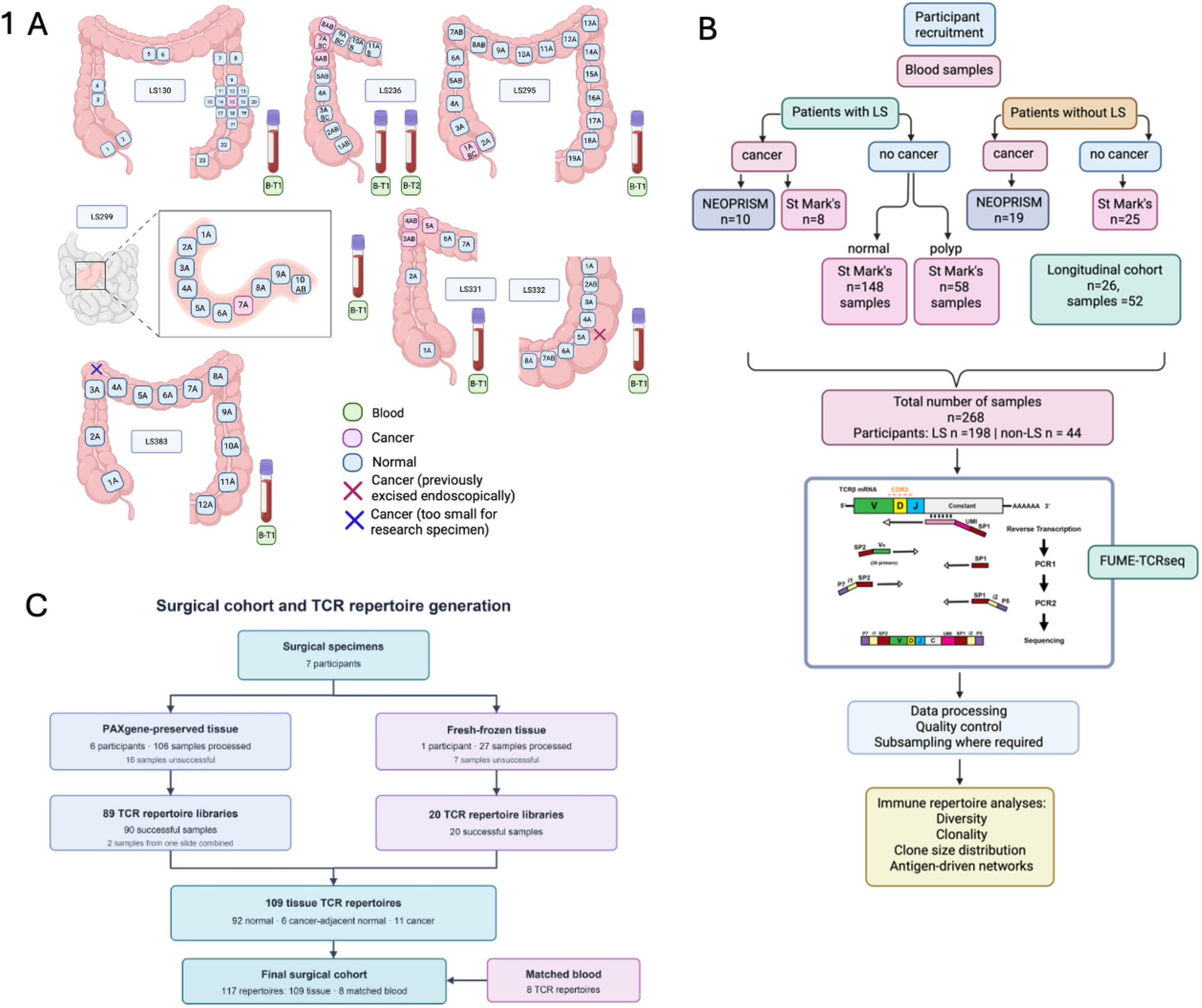
Study cohorts, spatial sampling and TCR repertoire generation. **(A)** Anatomical distribution of colorectal tissue and matched peripheral blood samples collected from seven participants with Lynch syndrome in the surgical cohort. Numbered labels identify individual tissue samples. Colours indicate normal tissue, cancer and blood. Crosses mark cancers that had previously been removed endoscopically or were too small to provide a research specimen. B-T1 and B-T2 denote blood collected at the first and second timepoints, respectively. The inset shows detailed sampling around the tumour in participant LS299. **(B)** Recruitment and classification of the peripheral blood cohort. Participants with and without Lynch syndrome were stratified by cancer status and recruitment source. Participants with Lynch syndrome but without cancer were further classified according to contemporaneous endoscopic findings as normal or polyp. The cohort comprised 268 blood samples from 198 participants with Lynch syndrome and 44 participants without Lynch syndrome. Twenty-six participants contributed paired longitudinal samples, giving 52 longitudinal samples. **(C)** Overview of the surgical cohort and generation of TCRβ repertoires. Multi-region surgical specimens from seven participants and peripheral blood samples were processed using the unique molecular identifier-based FUME-TCRseq workflow to identify TCRβ CDR3 sequences. **Abbreviations:** CDR3, complementarity-determining region 3; LS, Lynch syndrome; TCR, T-cell receptor; UMI, unique molecular identifier

Our cross-sectional blood cohort comprised 268 repertoires from 242 participants including 198 LS carriers (Figure 1D). Longitudinal analysis included 52 samples from 26 LS participants with intervals ranging from 10.2 to 53.4 months (Supplementary Figure 8A). Comparator cohorts included baseline peripheral blood samples from the NEOPRISM clinical trial comprising patients with stage IIb or stage III dMMR cancer^11^ who received neoadjuvant immunotherapy (n=29, n=10 had confirmed LS, the remainder sporadic dMMR, Figure 1D) and a non-LS healthy cohort n=25 collected from St Mark’s Hospital (Figure 1D). Of the LS blood samples collected, 78 patients had a history of an LS-associated cancer (defined as CRC, endometrial, ovarian, gastric, urothelial, sebaceous carcinoma). The germline pathogenic variant split was *MLH1* 34.3% (n=68), *MSH2* 32.8% (n= 65), *MSH6* 12.6% (n=25), *EPCAM* 2.0% (n=4), *PMS2* 11.6% (n=23), unknown 6.6% (n=13). Total unique clonotype counts for each sample type were as follows: blood 13,996 (IQR 9,398-21,006), normal tissue 3,125 (IQR 1,556-5,156), cancer tissue 1,472 (IQR 1,176-3,874) and polyp 393 (IQR 235-502).

### Peripheral blood repertoires reflect local immune responses across the Lynch syndrome bowel

To determine whether systemic immune repertoires reflect local immune activity within the LS bowel, we calculated pairwise Morisita-Horn overlap between patient-matched TCR repertoires. For formal inference, median overlap was calculated for each participant within each compartment comparison (representative heatmaps Figure 2C LS236 and LS295, all patients Figure 2E). We found that TCR repertoires were more similar within the same compartment ie. “normal”, “cancer” than across different compartments ie “cancer-normal”. The median participant-level Morisita overlap between multi-region cancer repertoires was 0.411, compared with 0.598 between normal bowel repertoires. These overlaps did not differ significantly (model estimated normal-normal minus cancer-cancer difference, 0.169; BH adjusted *P* = 0.169). By contrast, the median cancer-normal overlap was 0.121, which was significantly lower than both normal-normal overlap (estimated difference, 0.482; BH-adjusted *P* = 1.99 x 10^-6^) and cancer-cancer overlap (estimated difference, -0.313; BH-adjusted *P* = 0.011).

**Figure 2.**
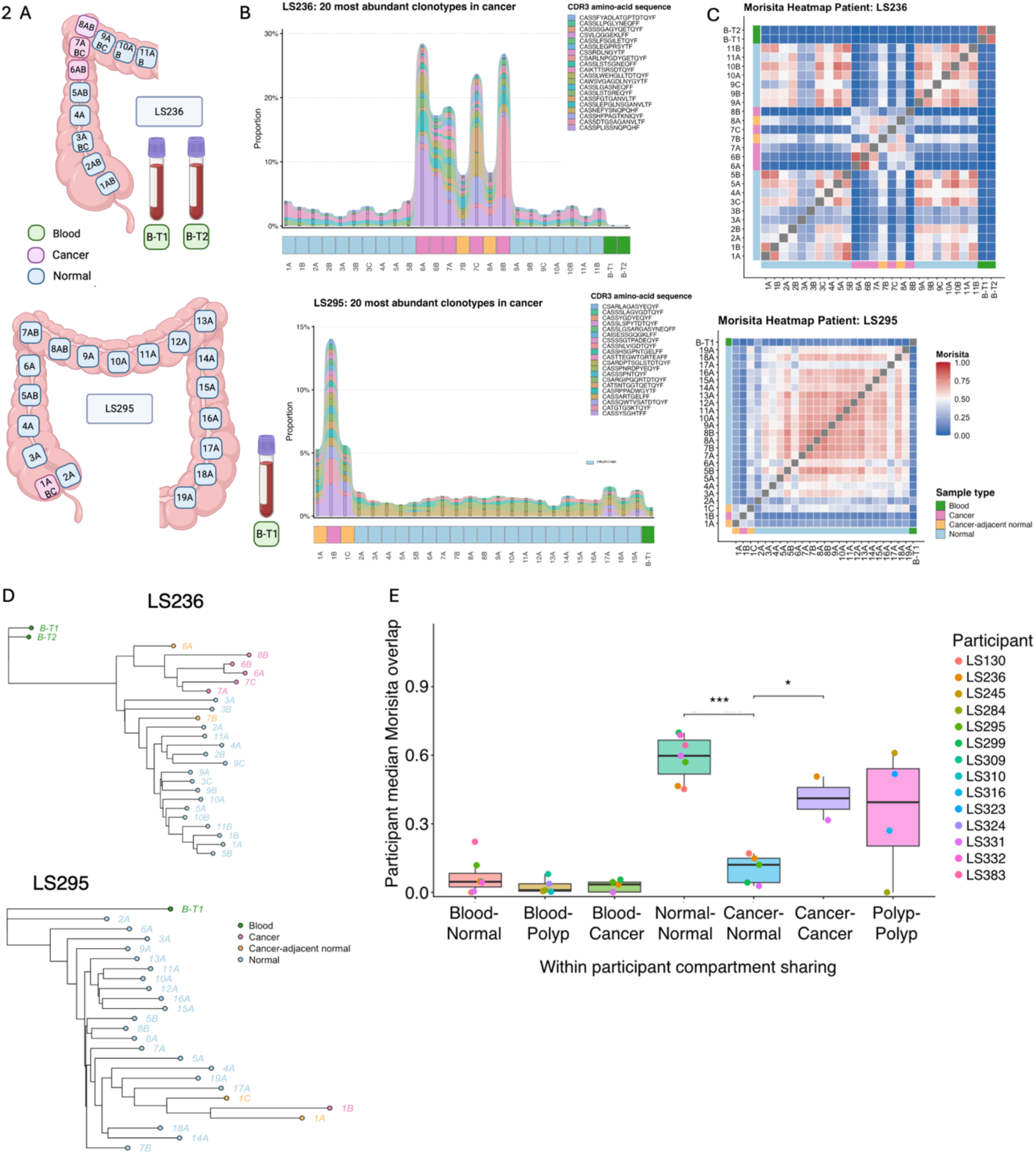
Spatial T-cell receptor repertoire architecture in two representative participants with Lynch syndrome. **(A)** Schematic sampling maps for LS236 (upper) and LS295 (lower), showing the anatomical locations of colorectal tissue samples and matched peripheral blood samples. Colours indicate normal tissue, cancer and blood. Sample identifiers correspond across panels. B-T1 and B-T2 denote blood collected at the first and second timepoints, respectively. **(B)** Distribution of the 20 most abundant cancer clonotypes across matched tissue and blood repertoires. Each coloured ribbon represents a unique CDR3 amino-acid sequence, and ribbon height shows its relative abundance within each sample. The annotation strip below each plot indicates sample type. **(C)** Pairwise Morisita–Horn overlap between TCR repertoires from the same participant. Values range from 0, indicating no repertoire overlap, to 1, indicating complete overlap. Sample-type annotations identify blood, cancer, cancer-adjacent normal and normal tissue. LS236 is shown above and LS295 below. **(D)** Neighbour-joining trees showing relationships between participant-matched T-cell receptor repertoires for LS236 (upper) and LS295 (lower). Trees were constructed from pairwise repertoire distances defined as 1 minus the Morisita–Horn overlap, calculated using CDR3 amino-acid clonotypes. Only samples containing more than 100 clonotypes were included. The LS236 tree was rooted on the first-timepoint blood sample, B-T1, and the LS295 tree was rooted on its eligible blood sample. Terminal nodes represent individual samples and are coloured by histology: blood, cancer, cancer-adjacent normal or normal tissue. Branch lengths represent repertoire dissimilarity within the neighbour-joining reconstruction and do not represent evolutionary time.**(E)** Within-participant T-cell receptor repertoire overlap across peripheral blood and colorectal tissue compartments. Pairwise Morisita–Horn overlap was calculated between participant-matched repertoires, and the median overlap for each participant within each compartment comparison is shown. Points represent participant-level medians and are coloured by participant. Boxplots show the median and interquartile range, with whiskers extending to 1.5 times the interquartile range. Normal–normal overlap was greater than cancer– normal overlap (BH-adjusted *P* = 1.99 × 10⁻⁶), and cancer–cancer overlap was greater than cancer–normal overlap (BH-adjusted *P* = 0.011). Normal–normal and cancer–cancer overlap did not differ significantly (BH-adjusted *P* = 0.169). \**P* < 0.05; \*\*\*\**P* < 0.001.

Blood–tissue repertoires had substantially lower overlap (median Morisita = 0.040; IQR 0.0004-0.055; n = 12 participants, Figure 2E and Supplementary Figure 1A-B). However, there was no evidence that overlap differed between blood-normal, blood-polyp and blood-cancer comparisons (all BH-adjusted *P* > 0.685). Circulating repertoires therefore captured a limited subset of the bowel repertoire, but the extent of this representation was not associated with tissue histology.

We compared repertoire overlap between samples from the same participant with overlap between different participants. Within-participant overlap was consistently greater than between-participant overlap across blood-tissue and tissue-tissue comparisons (Supplementary Figure 1B), confirming that the observed repertoire architecture was strongly participant specific.

Collectively, these findings demonstrate participant-specific and compartmentally structured TCR repertoire organisation in LS. Sharing of TCR clonotypes across normal and cancer tissue indicated that some tumour-associated T cells are not entirely tumour restricted, while the relatively limited blood-tissue overlap suggest that peripheral blood provides only a partial representation of the bowel immune repertoire.

### Circulating clonotype abundance predicts tissue detection

The probability that a circulating clonotype was detected in participant matched colorectal tissue increased markedly with its abundance in blood (Figure 3A). Relative to small clonotypes occupying 0.001-0.01% of the blood repertoire, the adjusted odds of tissue detection were approximately 7-fold greater for medium clonotypes (0.01-<0.1%), 57-fold greater for large clonotypes (0.1-<1%) and 188-fold greater for hyperexpanded clonotypes (≥1%; all *P* <2 x 10^-16^). Corresponding adjusted detection probabilities were 0.45%, 3.0%, 20.3% and 45.7%. After accounting for circulating abundance, participant and the number of tissue samples, overall detection did not differ between cancer and normal tissue (OR 1.09, 95% CI 0.76-1.58; *P* = 0.633). The abundance-by-histology interaction did not reach statistical significance (*P* = 0.052).

**Figure 3.**
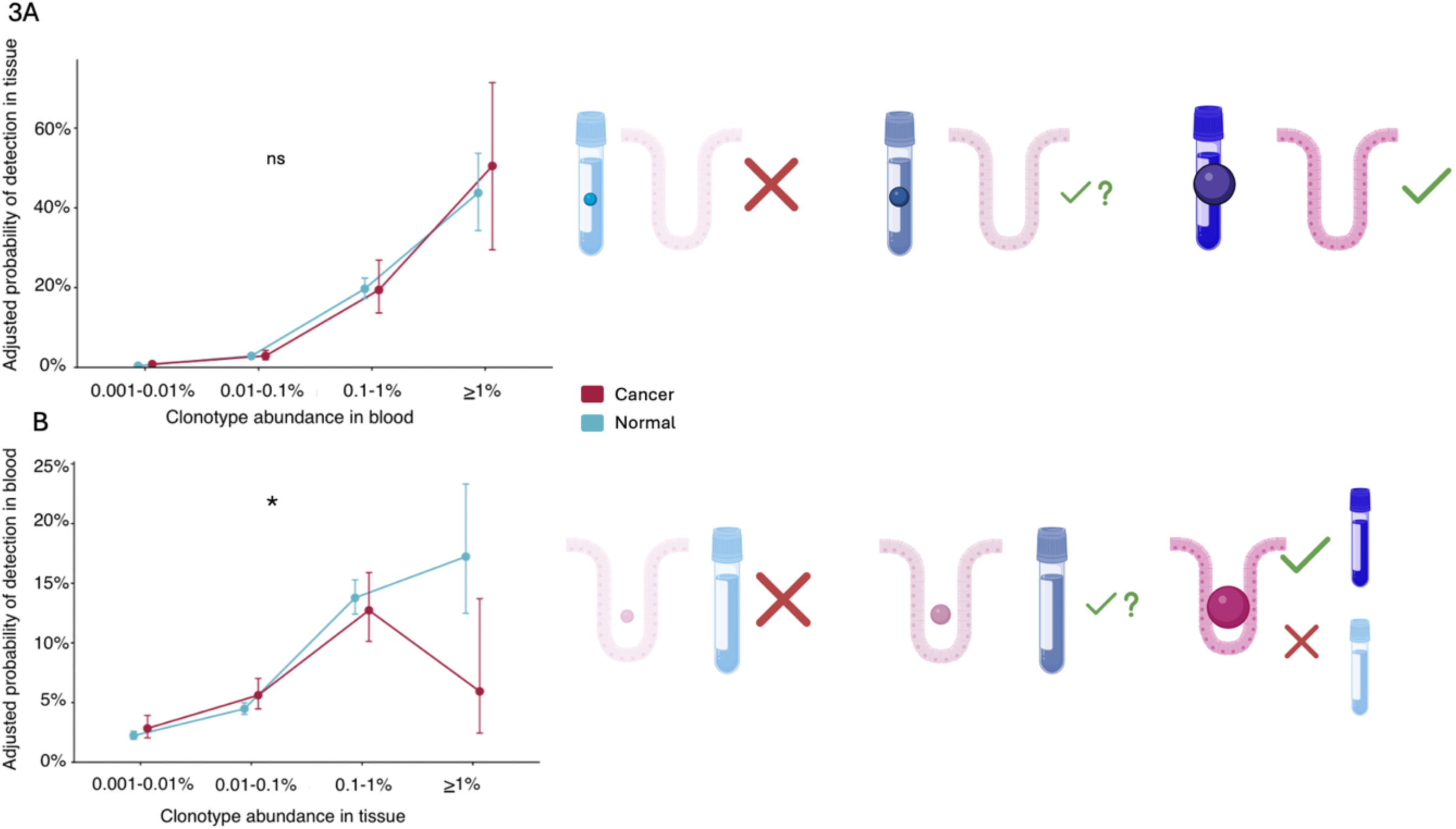
Abundance-dependent sharing of T-cell receptor clonotypes between peripheral blood and colorectal tissue. **(A)** Model-adjusted probability that a clonotype detected in peripheral blood was also detected in matched colorectal tissue, stratified by clonotype abundance in blood and tissue histology. The probability of tissue detection increased with clonotype abundance in blood. The association between abundance and detection did not differ significantly between cancer and normal tissue (abundance-by-histology interaction, *P* = 0.052). **(B)** Model-adjusted probability that a clonotype detected in colorectal tissue was also detected in matched peripheral blood, stratified by clonotype abundance in tissue and tissue histology. The association between abundance and blood detection differed between cancer and normal tissue (abundance-by-histology interaction, *P* = 0.002). In the highest abundance category, cancer-derived clonotypes had a lower predicted probability of detection in blood than clonotypes from normal tissue

In the reciprocal analysis, tissue clonotype abundance also predicted detection in matched peripheral blood, although this relationship differed according to tissue histology (abundance-by-histology interaction *P* = 0.002; Figure 3B). In normal tissue, the adjusted probability of blood detection increased from 2.23% among small clonotypes to 17.23% among hyperexpanded clonotypes. In cancer tissue, detection increased across the lower abundance categories but declined from 12.73% among large clonotypes to 5.94% among hyperexpanded clonotypes. This latter estimate was based on relatively few expanded cancer clonotypes and therefore should be interpreted cautiously.

To determine whether blood-tissue-shared clonotypes exhibited evidence of sequence convergence, sequence-similarity networks were constructed from the 3,000 most abundant circulating clonotypes in six participants. The median proportion belonging to a sequence similarity cluster increased from 9.0% among clonotypes not detected in tissue to 15.9% among tissue-detected clonotypes (paired Wilcoxon *P* = 0.036). After adjustment for blood abundance and participant level differences, tissue-detected clonotypes had approximately twice the odds of cluster membership (OR 1.96, 95% CI 1.75-2.19; *P* < 2 x 10^-16^). Blood abundance itself was not independently associated with cluster membership (OR per 10-fold increase 0.90, 95% CI 0.74-1.08; *P* = 0.247).

Tissue-detected clonotypes were also more extensively connected within circulating sequence-similarity networks: their expected network degree was 2.44 times greater than that of clonotypes not detected in tissue (IRR 2.44, 95% CI 2.16-2.72; *P* < 2 x 10^-16^). This association was independent of blood abundance, which was not associated with network degree (IRR per 10-fold increase 1.04, 95% 0CI 0.89-1.22; *P* = 0.641).

Collectively, these findings demonstrate that clonotype abundance is a major determinant of cross-compartment detection and that circulating clonotypes shared with colorectal tissue are preferentially embedded within sequence related networks, independently of their abundance.

### Convergent clonal architectures suggest recurrent systemic antigen-driven immune responses

To investigate whether cancer-associated T-cell responses in LS exhibit evidence of convergent antigen-driven expansion, we examined the spatial organisation and sequence-based clustering of the TCR clonotypes in each compartment (blood, normal tissue and cancer tissue). Most clonotypes were compartment-restricted, with the largest fractions detected only in blood, only in normal tissue or only in cancer.

Network-based clustering analysis, using string-kernel similarity matrices, identified large, interconnected clonotype structures (“meta-clonotypes”) shared across tissue and blood compartments within individual patients (Supplementary Figure 2A, LS236, LS130, LS295 and LS332). These networks included meta-clonotypes detected in blood, normal tissue and cancer tissue, as well as meta-clonotypes shared across these compartments. Network architecture varied markedly between participants: LS236 and LS295 contained numerous meta-clonotypes shared between normal and cancer tissue whereas LS130 was predominantly characterised by meta-clonotypes only found in normal bowel.

To evaluate cluster stability, we performed 100 bootstrap replicates. The mean proportion of hyperexpanded clonotypes was positively correlated with the mean size of the largest sequence similarity cluster, although this did not reach statistical significance (Spearman ρ = 0.90, *P* = 0.083, n = 5; Supplementary Figure 2B). Among the four participants with colonic cancers, tumour size was positively correlated with the mean proportion of cancer associated clusters (Spearman ρ = 1, *P* = 0.083; Supplementary Figure 2C). By contrast, the proportion of hyperexpanded clonotypes was not associated with the proportion of cancer-associated clusters (Spearman ρ = −0.1, *P* = 0.95).

### Peripheral blood TCR clustering distinguishes cancer-free LS carriers from non-LS controls

Analysis of expanded T-cell receptor (TCR) clustering architecture was performed across peripheral blood repertoires from LS and non-LS cohorts to investigate whether TCR repertoire organisation differed according to LS-status, cancer status, prior cancer history, or age (Figure 4A). We examined four metrics: the total number of clustered CDR3 sequences, maximum cluster size, number of clusters and clusters containing hyperexpanded clonotypes (Figure 4B).

**Figure 4.**
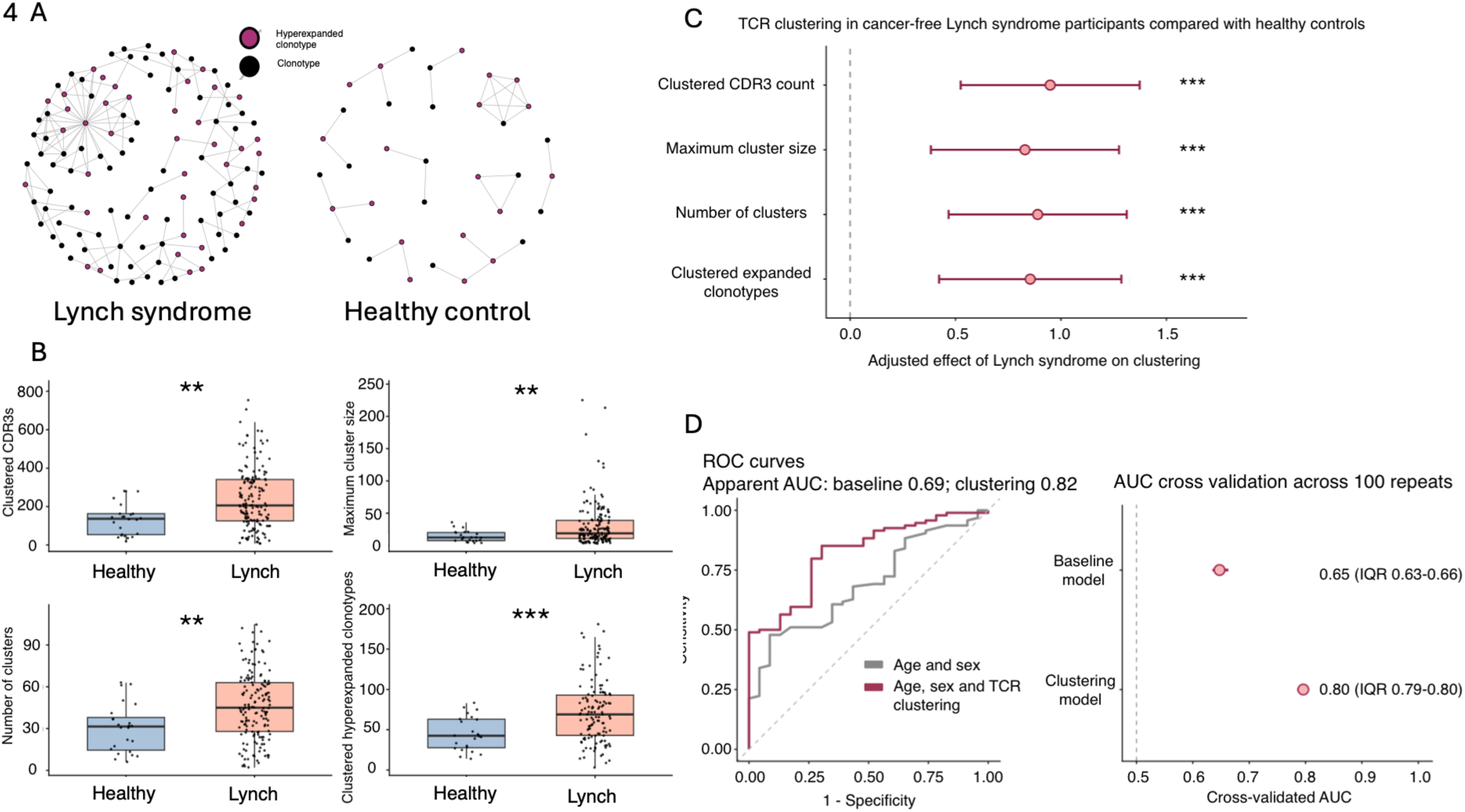
Systemic TCR sequence-similarity clustering distinguishes cancer-free participants with and without Lynch syndrome. **(A)** Representative TCR sequence-similarity networks generated from peripheral blood repertoires from a participant with Lynch syndrome (LS), showing extensive clustering, and a non-LS/healthy control, showing fewer and smaller connected structures. Each node represents a unique CDR3 amino-acid clonotype selected from the 3,000 most abundant clonotypes in that participant’s repertoire. Edges connect clonotypes with a normalised spectrum-kernel similarity score of at least 0.82, based on shared amino-acid triplets. Isolated clonotypes were excluded; only clonotypes belonging to a connected component containing at least two nodes are displayed. Network position and edge length are used solely for visualisation and do not represent biological distance. Quantitative clustering metrics were calculated separately. **(B)** Unadjusted distributions of clustered CDR3 count, maximum cluster size, number of clusters and clustered hyperexpanded clonotype count in cancer-free LS and non-LS participants. Points represent individual participants; boxplots show the median and interquartile range. *P* values were calculated using two-sided Wilcoxon rank-sum tests. **(C)** Adjusted differences in TCR clustering between cancer-free participants with LS and cancer-free non-LS controls. Points and error bars show adjusted coefficients (β) and 95% confidence intervals from separate multivariable linear regression models for four prespecified clustering metrics. Outcomes were log-transformed and standardised before modelling. Models were adjusted for age and sex; positive coefficients indicate greater clustering in LS. *P* values were adjusted across the four outcomes using the Benjamini–Hochberg false-discovery-rate procedure. **(D)** Receiver operating characteristic curves for a baseline logistic regression model containing age and sex and an extended model additionally containing standardised clustered CDR3 count. Apparent areas under the curve (AUCs) were 0.69 and 0.82, respectively. Internal validation using 100 repetitions of stratified five-fold cross-validation. Points show median cross-validated AUCs and horizontal lines indicate interquartile ranges: 0.65 (0.63–0.66) for the baseline model and 0.80 (0.79–0.80) for the extended model. Predictions for each fold were generated for participants excluded from model fitting.

Among cancer-free LS carriers there was greater TCR sequence-similarity clustering across all four metrics than cancer-free non-LS controls in adjusted comparisons: clustered CDR3 count (*P* = 0.001), maximum cluster size (*P* = 0.00046), number of clusters (*P* = 0.0022) and clustered hyperexpanded clonotype count (*P* = 0.00047; Figure 4B). These differences persisted after adjustment for age and sex. LS was associated with a higher clustered CDR3 count (standardised β = 0.95, 95% CI 0.52-1.37; BH-adjusted *P* < 0.0001), maximum cluster size (β = 0.83, 95% CI 0.38–1.28; P = 0.0004), number of clusters (β = 0.89, 95% CI 0.47–1.31; P = 0.0001) and clustered hyperexpanded clonotype count (β = 0.85, 95% CI 0.42–1.29; P = 0.0002; Figure 4C).

We sought to determine if TCR clustering metrics could identify LS individuals. We found that clustered CDR3 count provided discriminatory information beyond demographic characteristics. Adding clustered CDR3 count to a model containing age and sex increased apparent AUC from 0.692 to 0.821 (Figure 4D). Under 100 repetitions of repeated five-fold cross validation, the median AUC was 0.80 (IQR 0.79-0.80), compared with 0.65 (IQR 0.63-0.66) for age and sex alone (Figure 4D).

We assessed a previously published Vilar previvor-control classifier^10^, which was based on sharing of specific CDR3 amino-acid sequences. In the 121 participants with complete co-variate data, the Vilar score was not independently associated with LS after adjustment for age, sex and repertoire richness (OR per SD increase 1.02, 95% CI 0.64-1.69; *P* = 0.948), and did not improve model performance. The apparent AUC increased from 0.705 for age and sex alone to 0.826 after incorporating repertoire richness but remained 0.824 after addition of the Vilar score.

The predictive power of TCR clustering appeared to be associated primarily with LS rather than current cancer status. Comparisons across LS and non-LS participants with and without current cancer showed no significant cancer-associated differences within either cohort. Within LS, participants with any cancer history (prior or current cancer) appeared to exhibit reduced total number of clustered CDR3 sequences, total number of clusters, and maximum cluster size (4Figure 3A), however these differences were attenuated following adjustment for age, sex and pathogenic variant (Supplementary Figure 3B) with confidence intervals crossing the null and all BH-adjusted *P* = 0.0736.

Collectively, we found that systemic TCR sequence-similarity clustering metrics can identify cancer-free LS carriers. The results are consistent with altered immune repertoire organisation preceding overt malignancy.

### Longitudinal blood repertoires demonstrate stable global diversity alongside dynamic clonal turnover

We next sought to determine how stable the LS-associated clonotypes are over time. Longitudinal peripheral blood TCR repertoires from 26 patients undergoing surveillance endoscopy were examined. The median interval between paired blood samples was 24.1 months (range 10.2–53.4 months) (Figure 5A), consistent with UK surveillance practice.

**Figure 5.**
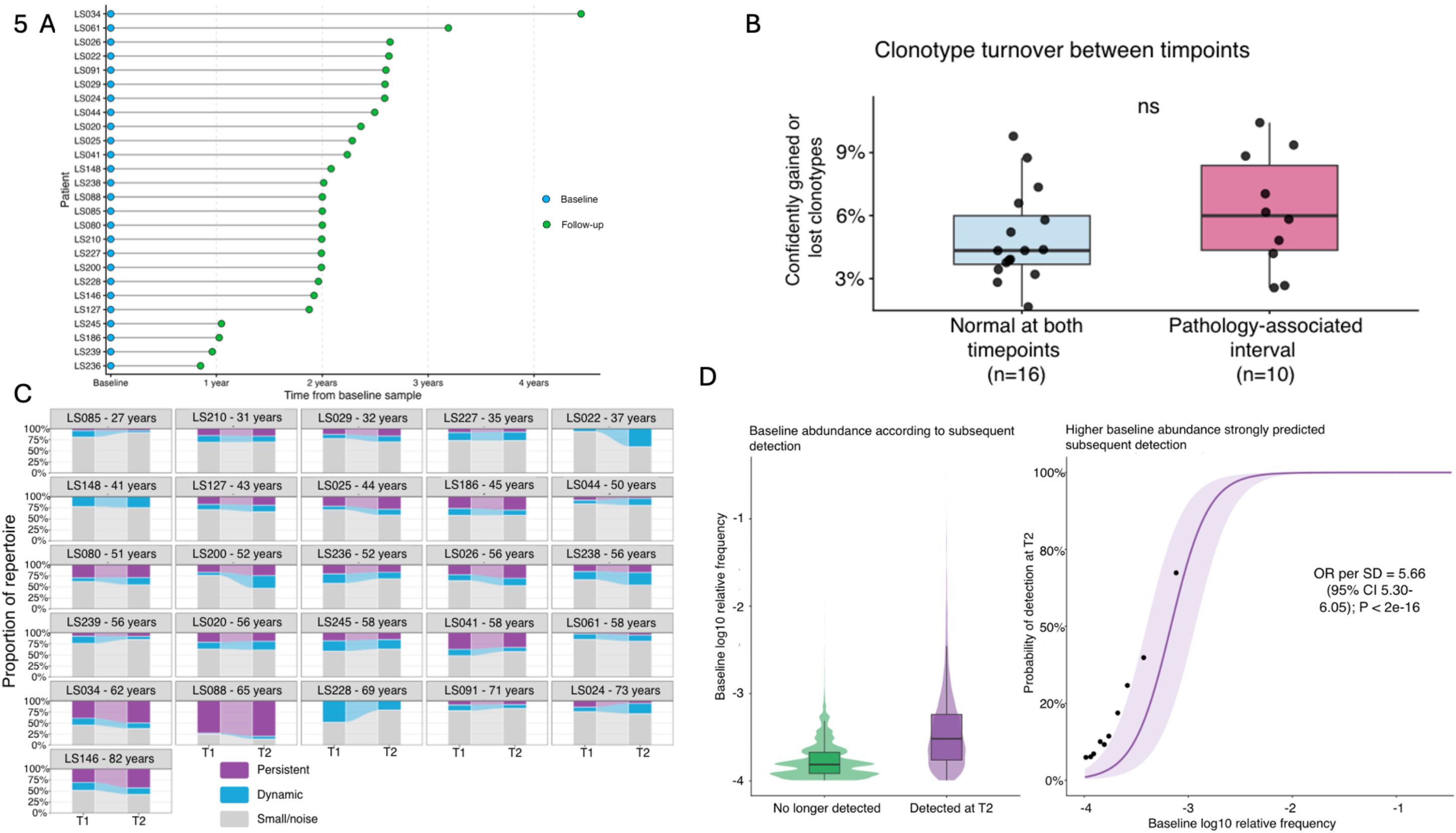
Longitudinal TCR repertoires show clonotype turnover against an abundance-dependent persistent repertoire backbone. **(A)** Whole-repertoire clonotype turnover in participants with normal findings at both timepoints and those with a pathology-associated interval. Turnover was defined as the number of confidently newly detected or no-longer-detected clonotypes divided by the total number of unique CDR3 sequences observed across either timepoint. Points represent individual participants; boxplots show the median and interquartile range. Turnover was numerically higher during pathology-associated intervals than during normal–normal intervals, but the difference was not statistically significant (median 5.99% versus 4.33%; Wilcoxon rank-sum test, *P* = 0.257). **(B)** Longitudinal peripheral blood TCRβ repertoire composition ordered by participant age. Participants are arranged from youngest to oldest, with age calculated at the midpoint between T1 and T2. For each participant, bars show the proportions of the repertoire occupied by three longitudinal clonotype compartments. Persistent clonotypes (purple) were confidently detected at both timepoints, including clonotypes that expanded or contracted. The gained-or-lost compartment (blue) comprised clonotypes confidently detected at only one timepoint, while the remaining repertoire (grey) comprised clonotypes that did not meet either definition. Connecting bands show changes in the cumulative relative abundance of each compartment and do not represent trajectories of individual clonotypes. Persistent repertoire burden showed a modest, non-significant association with age (Spearman’s ρ = 0.292; BH-adjusted *P* = 0.198). **(C–D)** Association between baseline clonotype abundance and subsequent detection, including 20,155 CDR3 amino-acid clonotypes confidently detected at T1 across 26 participants. **(C)** Distribution of baseline log₁₀ relative frequencies for clonotypes subsequently detected at T2 and those no longer detected. Violin plots show abundance distributions and embedded boxplots show the median and interquartile range. **(D)** Modelled probability of clonotype detection at T2 according to baseline log₁₀ relative frequency. The line shows population-level predictions from a mixed-effects logistic regression model with a participant-level random intercept; the shaded band indicates the 95% confidence interval. Points show observed detection proportions within deciles of baseline abundance. Each one-standard-deviation increase in baseline abundance was associated with 5.66-fold greater odds of subsequent detection (95% CI 5.30–6.05; *P* < 2 × 10⁻¹⁶). This relationship may reflect both biological persistence of abundant clonotypes and their greater probability of being captured during repertoire sampling.

Clonotype level turnover was evident despite broadly stable global repertoire architecture. Clonotypes newly detected or no longer detected between timepoints represented a median of 4.60% unique clonotypes observed across each sample pair. Turnover was numerically greater during pathology-associated intervals than during intervals with normal findings at both timepoints (5.99% versus 4.33%), but this difference was not statistically significant (*P* = 0.257; Figure 5B). Clonal turnover demonstrated no significant correlation with sampling interval (Spearman ρ = 0.18, *P* = 0.46), suggesting stability of the repertoire was not associated with time.

The relative contributions of newly detected and no longer detected clonotypes varied between participants and clinical transitions (Supplementary Figure 4A-B). However, transition groups were small and unequal (the normal-polyp and polyp-polyp groups contained four participants, polyp-normal and cancer-normal groups each contained only one), therefore statistical power was limited.

The proportion of persistent clonotypes did not change significantly between timepoints (*P* = 0.196, Figure 5B). Among 20,155 clonotypes confidently detected at baseline across 26 participants, higher baseline abundance strongly predicted longitudinal persistence. Each 1-SD increase in baseline log_10_ relative frequency was associated with 5.66-fold higher odds of detecting the same clonotype at follow up (95% CI 5.30-6.05; *P* < 2 x 10^-16^; Figure 5D). This highlights that our quantification of persistent circulating clonotypes is likely a function of biological abundance and the greater probability of resampling abundant clonotypes.

Consistent with this qualification, modelling of persistent-clonotype abundance changes suggested that threshold crossing arising from sampling variation could account for approximately 51.1% of observed clonotype losses (Spearman ρ = 0.41; *P* = 0.036; Supplementary Figure 5A). Therefore, sampling differences only account for half of apparent change, suggesting that the remainder could be biologically driven.

Global repertoire diversity was also broadly conserved. After adjustment for total TCR count, Shannon entropy was not different by timepoint (*P* = 0.445), interval group (*P* = 0.373), or timepoint-by-interval group interaction (*P* = 0.431).

Collectively, these findings demonstrate clonotype-level turnover against a background of stable global TCR diversity in peripheral blood, with the most abundant clonotypes generally persisting through years.

## Discussion

In this study, we performed integrated spatial and longitudinal profiling of TCR repertoires across tumour, normal bowel, polyp and peripheral blood compartments in LS. Repertoires were strongly participant specific and with clear compartmentalisation (yet substantial clonotype sharing) between tumour and normal regions. Blood–tissue overlap was substantially lower, although abundant circulating clonotypes were more likely to be detected in colorectal tissue than rare clonotypes, indicating that local immune activity is partially represented systemically.

Cancer-free LS carriers exhibited greater circulating TCR sequence similarity-based clustering than cancer-free non-LS controls, independent of age and sex. This suggests that systemic immune repertoire alterations are a feature of LS, even before malignancy. We have shown in our cohort the potential power of repertoire features to identify LS carriers, although prospective studies are required to determine whether they predict subsequent neoplasia or cancer.

Longitudinally, global diversity and proportion of persistent clonotypes remained broadly stable despite ongoing clonotype gain and loss. Persistence was strongly abundance-dependent, supporting a durable backbone of higher-frequency circulating clonotypes. However, we did not identify any overt repertoire changes that consistently corresponded to contemporaneous colorectal neoplasia, indicating that any clinically useful TCR-based biomarker of cancer risk may require longitudinal, within-person measurements rather than a single global repertoire metric.

Sequence-similarity networks identified heterogenous but spatially distributed meta-clonotype architectures within individual participants. Meta-clonotypes spanning blood and colorectal tissue were consistent with coordinated antigen-driven immune responses across anatomical compartments. Exploratory associations with clonal expansion and tumour size provide a rationale for validation in larger cohorts.

The limited transferability of the Vilar model^10^ may reflect differences in cohort composition, HLA background, environmental exposure or sequencing methodology and suggests that exact public clonotype signatures may be cohort dependent. Higher order repertoire features, such as sequence similarity architecture, may prove more transferable, although we note that this requires independent external validation.

An association between observed repertoire structures and recurrent dMMR-derived frameshift neoantigens remains biologically plausible. However, we did not perform functional validation of antigen specificity in this study. Larger prospective cohorts integrating neoantigen prediction, HLA genotype and functional T-cell assays will be required to establish whether the presence of LS-associated meta-clonotypes can support cancer-risk prediction, or if their cognate neoantigen(s) represents a target for immunoprevention.

Together these findings position the circulating TCR repertoire as a measurable record of immune surveillance in LS and a candidate platform for future risk stratification and vaccine development.

## Methods

The methodological approach was designed to enable integrated analysis of local and systemic T-cell responses in Lynch syndrome pathogenic variant carriers.

### Ethical considerations

Human samples were collected and analysed under appropriate institutional ethical approvals and governance procedures. Samples collected at St Mark’s Hospital and Academic Institute (LS patients and non-LS healthy controls) were obtained through the LynC study (IRAS 276293), with written informed consent obtained for prospectively recruited participants. Additional samples from external cohorts, including NEOPRISM and comparator datasets, were analysed under independent ethically approved study protocols and institutional data-sharing agreements.

### Sample collection and processing

Peripheral blood and tissue samples were collected from individuals undergoing surveillance endoscopy or surgical resection.

Peripheral blood samples collected during surveillance were obtained in EDTA tubes. At the time of surgical resection (colectomy), additional blood samples were collected in multiple tube types, including EDTA tubes (BD Vacutainer), Tempus™ RNA tubes (Applied Biosystems™, ThermoFisher Scientific, United Kingdom, Catalogue number: 4342792) and Streck DNA tubes (Alpha Laboratories, United Kingdom, Catalogue number: 218997), to enable downstream nucleic acid analyses.

Tissue samples obtained at endoscopy were embedded in optimal cutting temperature (OCT) compound within cryomoulds and frozen at −80°C without prior snap freezing. For storage efficiency, frozen samples were subsequently transferred to cryovials and stored at −80°C until further processing.

Surgical resection specimens were processed using multiple preservation approaches to support downstream analyses. Sections of tissue were fixed in PAXgene tissue fixative (PAXgene Tissue FIX container, Qiagen), followed by transfer to stabilisation solution (PAXgene Tissue STABILIZER concentrate Qiagen) after approximately 18 hours, according to the manufacturer’s protocol. Additional tissue samples were placed in RPMI-based transport media for immediate downstream applications where required. Parallel tissue samples were also snap-frozen directly in cryovials without embedding for RNA-based analyses.

This multi-modal sampling strategy enabled parallel assessment of tissue-resident and circulating immune responses across a range of preservation conditions.

### FUME TCR sequencing and library preparation

T-cell receptor (TCR) β-chain libraries were prepared using a multiplex PCR-based approach (FUME-TCRseq), designed to amplify the hypervariable complementarity-determining region 3 (CDR3) from total RNA. This method incorporates unique molecular identifiers (UMIs) during reverse transcription to enable accurate quantification of clonotypes. This was performed as previously described in paper by Baker et al. Sequencing was performed on the Illumina MiniSeq platform using a mid- or high-output flow cell or on the NovoSeq platform. Paired- end sequencing was carried out with read lengths of 151 base pairs and dual 8 bp index reads. Data acquisition and run monitoring were performed using Illumina BaseSpace software. Sequencing depth was sufficient to capture the breadth and clonal structure of the TCR repertoire across samples.

### TCR sequence processing

Raw sequencing data were demultiplexed and converted to FASTQ format using Illumina BaseSpace.

TCR sequences were processed using Decombinator^12^, which identifies V and J gene usage and extracts complementarity-determining region 3 (CDR3) sequences. Unique molecular identifiers (UMIs) incorporated during reverse transcription were used to collapse PCR duplicates, improving quantitative accuracy of clonotype frequencies. Reads were filtered to remove low-quality sequences and non-productive rearrangements where appropriate. Repertoires were quality-controlled by checking sample–metadata concordance and calculating the number of unique clonotypes per sample. Samples with ≤100 unique clonotypes were excluded. The number of samples passing QC and the median clonotype count (IQR) were summarised by tissue compartment.

### TCR repertoire analysis

Processed T-cell receptor (TCR) sequencing data were analysed to characterise features of antigen-driven immune responses across tissue and peripheral blood compartments. Analyses focused on quantifying clonal expansion, repertoire diversity, sequence similarity and overlap between samples with the aim of identifying signatures associated with Lynch syndrome and cancer status. Clonotype definition, TCR sequences were grouped into clonotypes based on identical exact match complementarity determining region 3 (CDR3) amino acid sequences. Each clonotype was assigned a frequency based on it’s relative abundance within the repertoire. Analysis was performed using the Immunarch package in R and custom pipelines. Repertoires were downsampled to uniform sequencing depth, where appropriate, using custom random subsampling pipelines implemented in Python.

### Cluster filtering and classification

Shared clonotypes were defined as identical CDR3 amino acid sequences detected across two or more anatomical or temporal compartments within an individual patient. Clonotypes identified within water-control libraries were removed from all downstream analyses prior to repertoire overlap and clustering analyses. Compartment sharing was subsequently summarised using overlap analyses.

For network-based clustering analyses, repertoires were restricted to the 3000 most abundant clonotypes within each sample to reduce noise from low-frequency sequences and enable comparison of dominant repertoire architecture across samples. Cancer-associated clusters were defined as sequence-similarity clusters seeded by clonotypes identified within tumour tissue. These clusters were subsequently tracked across matched blood and histologically normal bowel compartments to assess spatial conservation of tumour-associated immune responses.

For compartment-level repertoire analyses, hyperexpanded clonotypes were defined as clonotypes within the upper 1% of relative abundance within each repertoire. Summary metrics including clustered CDR3 burden, largest cluster size, and hyperexpanded repertoire proportion were subsequently calculated for each sample.

### Clustering similarity analysis

To identify groups of related TCR sequences potentially recognising shared or structurally related antigens, pairwise CDR3 amino acid sequence similarity was calculated using a spectrum string kernel^13^. Each CDR3 sequence was represented by its constituent amino acid k-mers, and pairwise similarity was computed from the inner product of these k-mer feature vectors. Kernel values were normalised to account for differences in CDR3 length, generating a similarity score between 0 and 1 for each pair of clonotypes.

A clonotype similarity network was then constructed for each sample, in which nodes represented unique CDR3 amino acid clonotypes and edges connected clonotypes with normalised similarity above a predefined threshold. A threshold of 0.75 was selected to identify highly similar CDR3 sequences within different compartments while excluding weak sequence relationships, within blood alone a more stringent threshold of 0.82 was selected. Connected components within this graph were defined as TCR sequence clusters, representing putative antigen-driven clonotype families. For each sample, cluster size was defined as the number of clonotypes within a connected component, and cluster abundance was calculated as the summed relative frequency of all clonotypes within that cluster.

### Bootstrap resampling analysis

To account for unequal compartment representation and sampling heterogeneity between patients, bootstrap resampling was performed across the spatial tissue cohort. For each iteration, a single blood, tumour and histologically normal tissue sample were randomly selected per patient, and clustering analyses repeated. This process was performed across 100 independent iterations, generating patient-level summary metrics including cancer-associated cluster abundance, hyperexpanded clonotype burden and largest cluster size. Correlation analyses were subsequently performed using patient-level mean values across bootstrap iterations.

Statistical and repertoire analyses were performed in R (version 4.5.2) using immunarch, igraph, kernlab, tidyverse and associated visualisation and mixed-effects modelling packages.

### Abundance-dependent blood-tissue clonotype sharing

Clonotype detection in matched blood and colorectal tissue was assessed bidirectionally using identical CDR3 amino acid sequences. Source-compartment abundance was classified using immunarch thresholds. Detection was modelled using beta-binomial mixed-effects regression, accounting for abundance category, tissue histology and participant, with a sample-level random intercept. Abundance by histology interactions were evaluated, and adjusted detection probabilities with 95% confidence intervals were calculated.

### Longitudinal clonotype tracking and persistence

Clonotypes were matched between paired blood samples (T1 and T2) using identical CDR3 amino acid sequences. Confident detection required ≥4 UMI-collapsed counts and >0.01% relative frequency. Clonotypes were classified as persistent, newly detected or no longer detected according to detection at both timepoints, T2 only or T1 only, respectively. Persistent clonotypes were further classified as expanded or contracted using log_2_ fold change thresholds, remaining clonotypes formed a residual category. Turnover was defined as newly detected plus no longer detected clonotypes divided by all unique CDR3 sequences observed across either timepoint. Compartment abundance was calculated by summing constituent clonotype frequencies. Turnover was compared between normal-normal and pathology associated intervals using Wilcoxon rank sum tests and correlated with sampling interval using Spearman’s correlation.

Among clonotypes confidently detected at T1, subsequent detection was modelled using mixed-effects logistic regression with standardised log_10_ baseline frequency and a participant random intercept. Odds ratios were reported per standard deviation increase, alongside predicted probabilities and 95% confidence intervals.

### Custom code

Custom scripts are available at https://github.com/pebbe-code/gedy-lab-tcr.git

## Supporting information

Blood demographic data

## Acknowledgements

PE was supported by the Tom Smith Fellowship, funded by 40tude. The LynC study was also supported by 40tude. The authors thank the Core Genomics Facility and Histopathology Services at The Institute of Cancer Research for technical support. We also thank the surgical and histopathology teams at Northwick Park Hospital and Central Middlesex Hospital for their assistance with sample collection and processing. We are grateful to James Reading, Petros Fessas and Amber Rogers at University College London for helpful discussions and advice. KB was funded by the Swiss National Science Foundation (P500PM_217647/1).

## Contributions

PE: Conceptualisation, Methodology, Investigation, Formal analysis, Data curation, Writing – original draft, Writing – review and editing. LS: Conceptualisation, Methodology, Investigation. TR: Software, Data curation. YJ: Resources, Investigation. GN: Investigation. CS: Investigation. SY: Investigation. HG: Data curation, Formal analysis. ES: Data curation, Formal analysis. KB: Investigation, Resources. SV: Investigation, Data curation. AM: Investigation, Resources. JF: Investigation. BC: Software, Data curation. KS: Resources, Investigation. AMB: Conceptualisation, Methodology, Investigation, Formal analysis, Data curation, Writing – review and editing. KM: Conceptualisation, Methodology, Investigation, Writing – review and editing. TG: Conceptualisation, Supervision, Writing – review and editing.

---

**Supplementary Figure 1.**
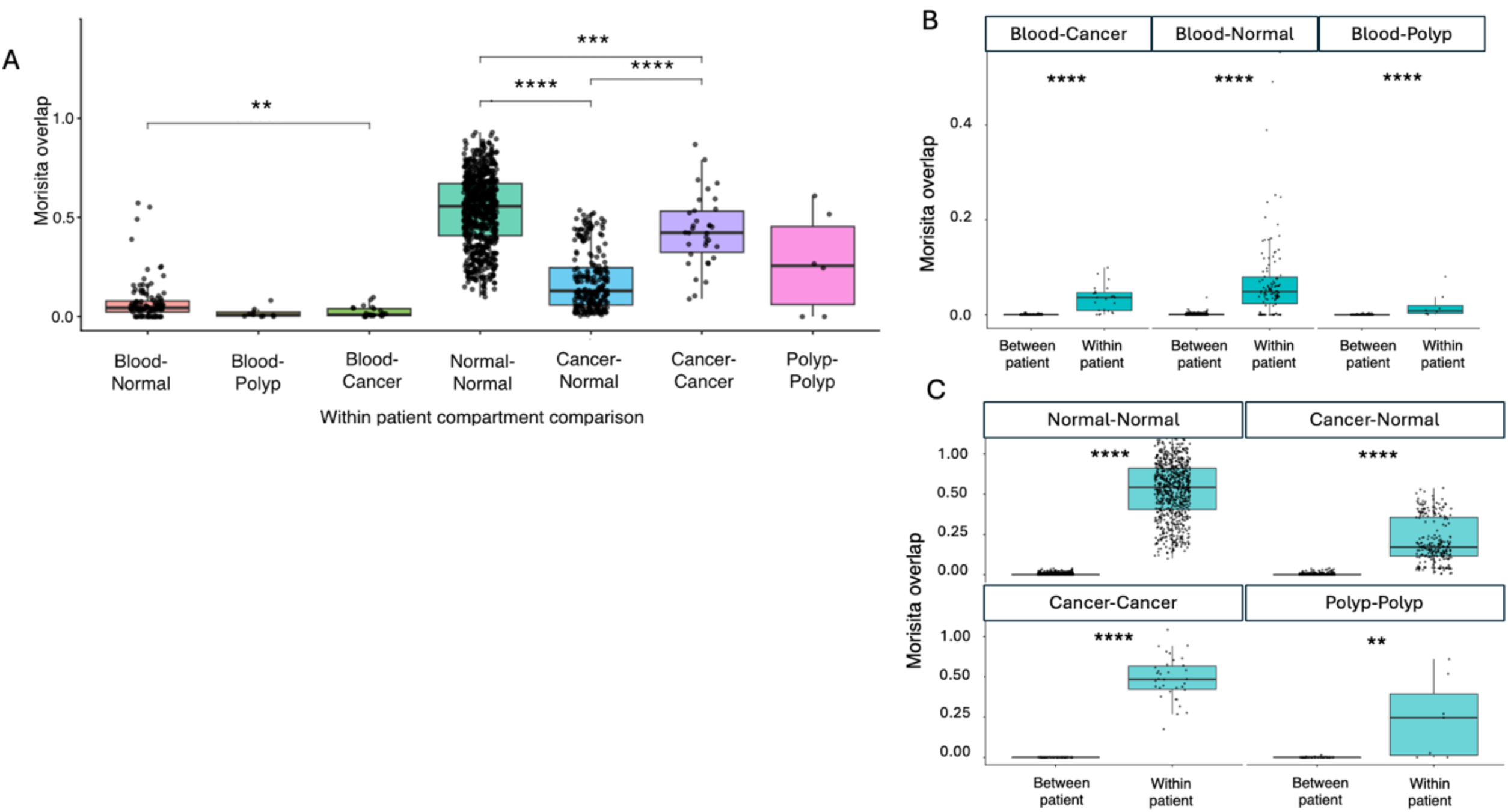
Pair-level TCR repertoire overlap across compartments and within versus between participants. **(A)** Pairwise Morisita–Horn overlap between peripheral blood and colorectal tissue compartments and between samples from the same tissue compartment. Each point represents one within-participant sample-pair comparison; boxplots show the median and interquartile range. Repertoire overlap differed significantly among compartment comparisons (Kruskal–Wallis test, *P* < 2.2 × 10⁻¹⁶); selected post-hoc pairwise *P* values are shown above the brackets. **(B)** Within-participant and between-participant overlap for blood–cancer, blood–normal and blood–polyp sample pairs. **(C)** Within-participant and between-participant overlap for normal–normal, cancer–normal, cancer–cancer and polyp–polyp sample pairs. In (B) and (C), each point represents one pairwise repertoire comparison, boxplots show the median and interquartile range, and *P* values were calculated using two-sided Wilcoxon rank-sum tests. Because individual samples contributed to multiple pairwise comparisons, the results in all panels are interpreted descriptively; participant-level mixed-effects analysis is presented in Figure 2A. The Morisita–Horn index ranges from 0, indicating no repertoire overlap, to 1, indicating identical repertoires.

**Supplementary Figure 2.**
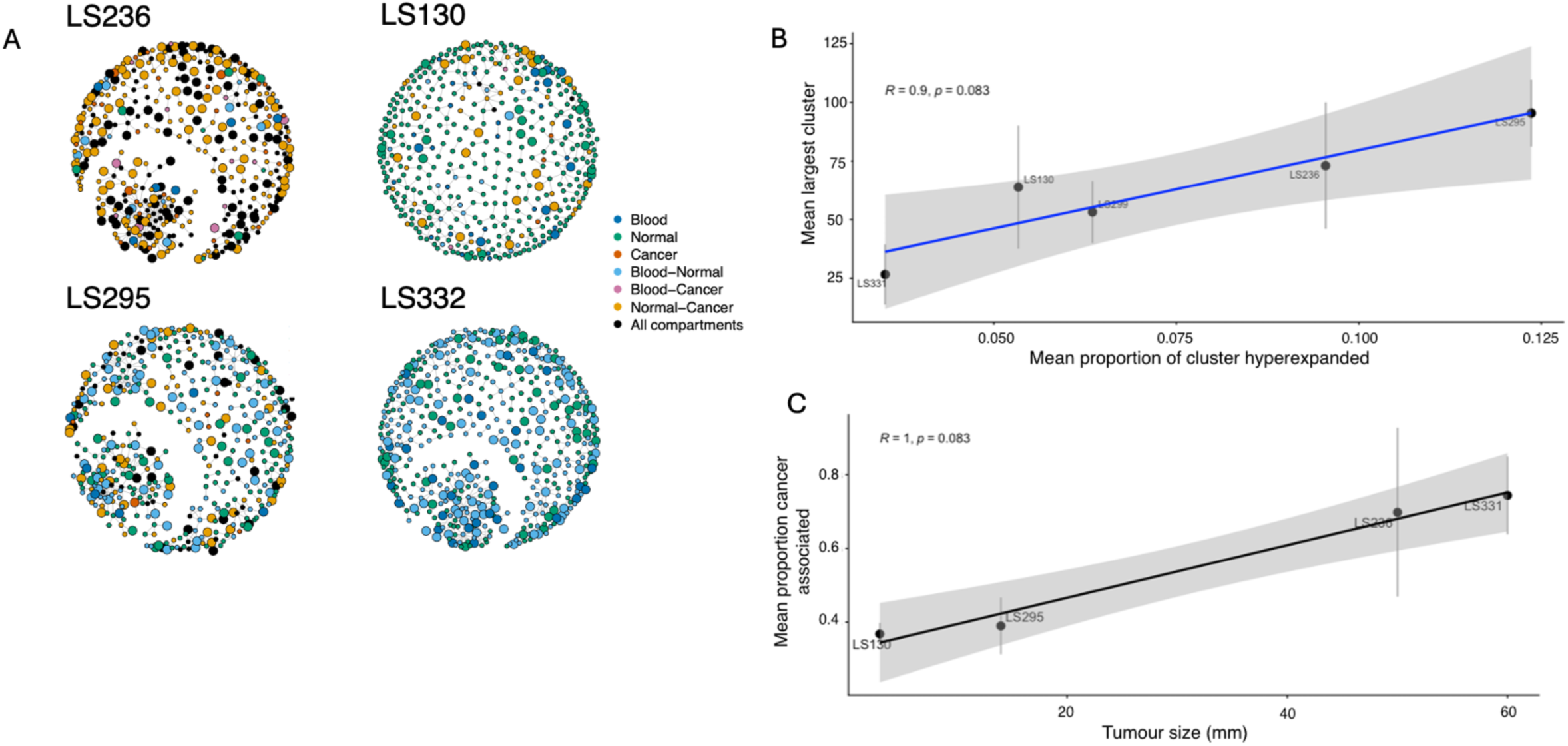
Multi-compartment TCR sequence-similarity networks and exploratory associations with clonal expansion and tumour size. **(A)** Representative within-participant TCR sequence-similarity networks for LS236, LS130, LS295 and LS332, respectively. Each node represents a unique CDR3 amino-acid clonotype and edges connect clonotypes with a normalised spectrum-kernel similarity score of at least 0.75, based on shared amino-acid triplets. Node size reflects clonotype abundance, while colour indicates the compartment or combination of compartments in which the clonotype was detected: blood, histologically normal tissue, cancer, blood–normal, blood–cancer, normal–cancer or all three compartments. Network position and edge length are used for visualisation and do not represent biological or anatomical distance. **(B)** Proportion largest cluster and proportion hyperexpanded. Relationship between the mean proportion of hyperexpanded clonotypes and mean largest sequence-similarity cluster size across five participants. The fitted line and shaded band show the estimated trend and 95% confidence interval. A positive correlation was observed but did not reach statistical significance (Spearman’s ρ = 0.90, *P* = 0.083; *n* = 5). **(C)** Proportion of cancer-associated clusters vs tumour size (colon only, n= 100 replicates). Relationship between tumour size and the mean proportion of clusters containing cancer-associated clonotypes among four participants with colonic cancers. Participant-level estimates were derived across 100 replicate networks; vertical error bars indicate variability across replicates. Tumour size correlated positively with the proportion of cancer-associated clusters, although the analysis was not statistically significant (Spearman’s ρ = 1.00, *P* = 0.083; *n* = 4). Given the small participant numbers, the associations in **B-C** should be considered hypothesis-generating.

**Supplementary Figure 3.**
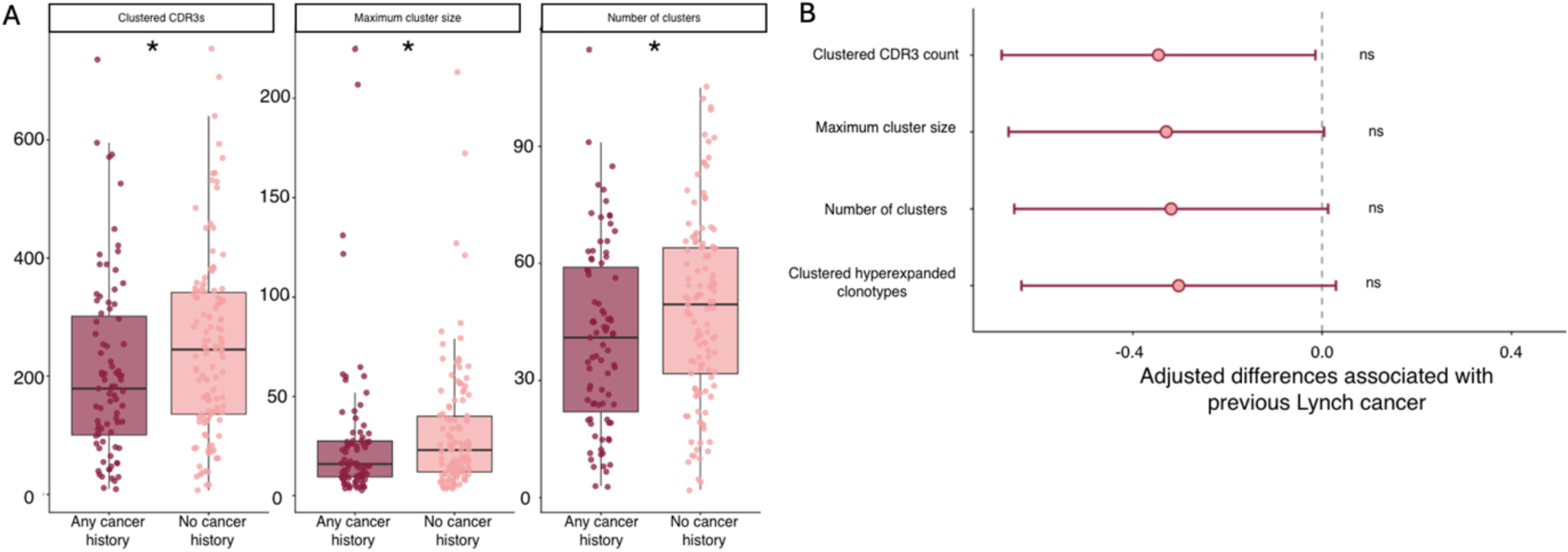
TCR clustering does not discriminate previous cancer history. **(A)** Unadjusted clustering metrics in LS carriers with any previous cancer history and those without a cancer history. Outcomes comprised clustered CDR3 count, maximum cluster size and number of sequence-similarity clusters. One blood repertoire was included per participant. Participants without a cancer history had higher values for all three metrics (Wilcoxon rank-sum *P* = 0.030, 0.031 and 0.026, respectively). Points represent individual participants; boxplots show the median and interquartile range. **(B)** Adjusted associations between previous LS-associated cancer and four TCR clustering metrics. Points show standardised regression coefficients and horizontal bars indicate 95% confidence intervals for participants with versus without previous LS-associated cancer. Models were adjusted for age, sex and pathogenic variant. Negative estimates indicate lower clustering among participants with a previous cancer. *P* values were corrected across the four outcomes using the Benjamini–Hochberg procedure. Although all estimates were negative, none remained statistically significant following correction (BH-adjusted *P* = 0.0736 for each outcome).

**Supplementary Figure 4.**
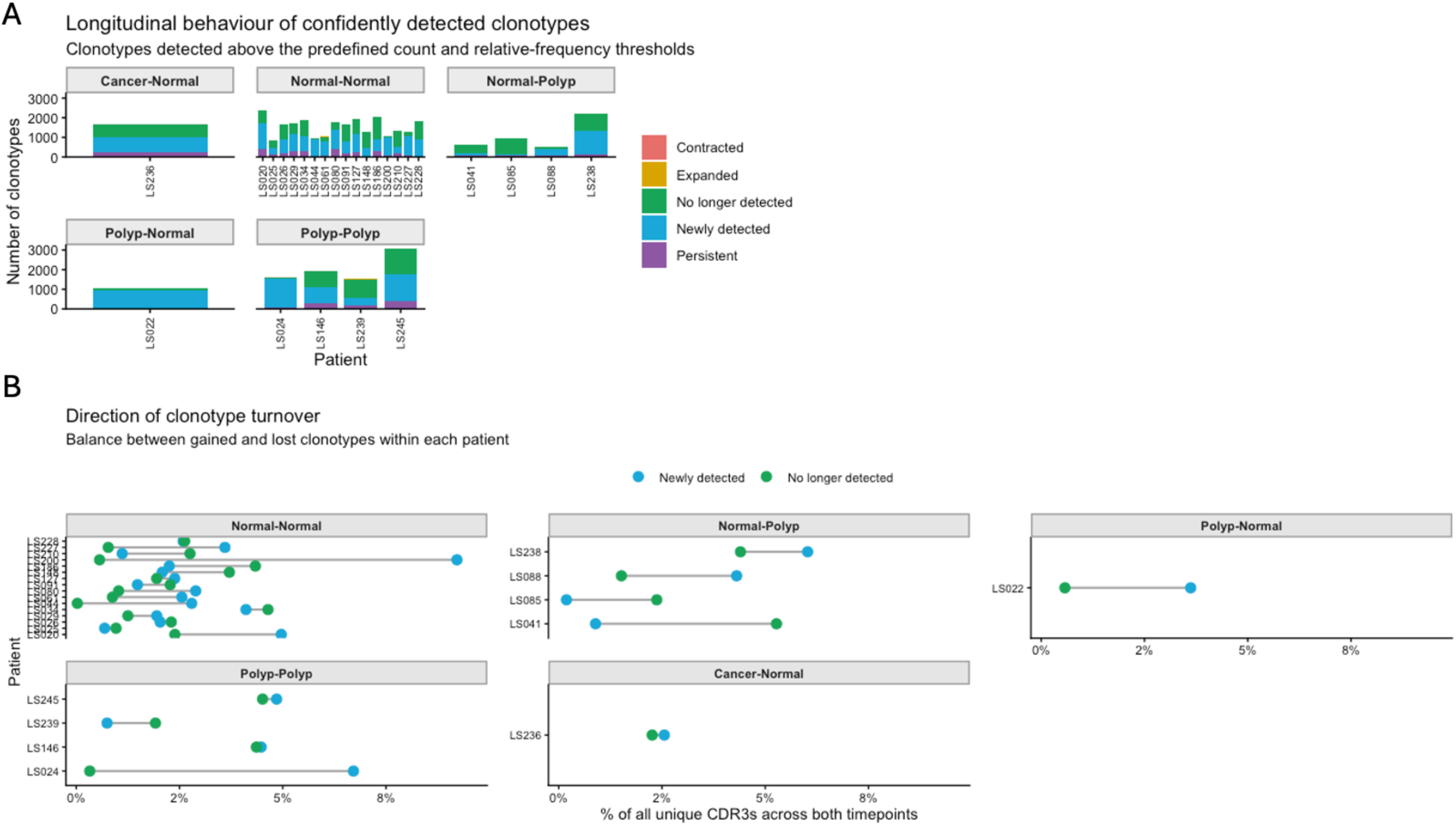
Longitudinal behaviour and direction of turnover of confidently detected circulating TCRβ clonotypes. Confident detection required at least three duplicate counts and a relative frequency of at least 0.01% at one or both timepoints. **(A)** Number of clonotypes within each longitudinal category for every participant, stratified by clinical transition between T1 and T2. Stacked bars show clonotypes classified as persistent, newly detected, no longer detected, expanded or contracted. **(B)** Participant-level balance between newly detected and no-longer-detected clonotypes, stratified by clinical transition. Values are expressed as percentages of all unique CDR3 amino-acid sequences observed across either timepoint. Blue and green points show newly detected and no-longer-detected clonotypes, respectively, and lines connect values from the same participan

**Supplementary Figure 5.**
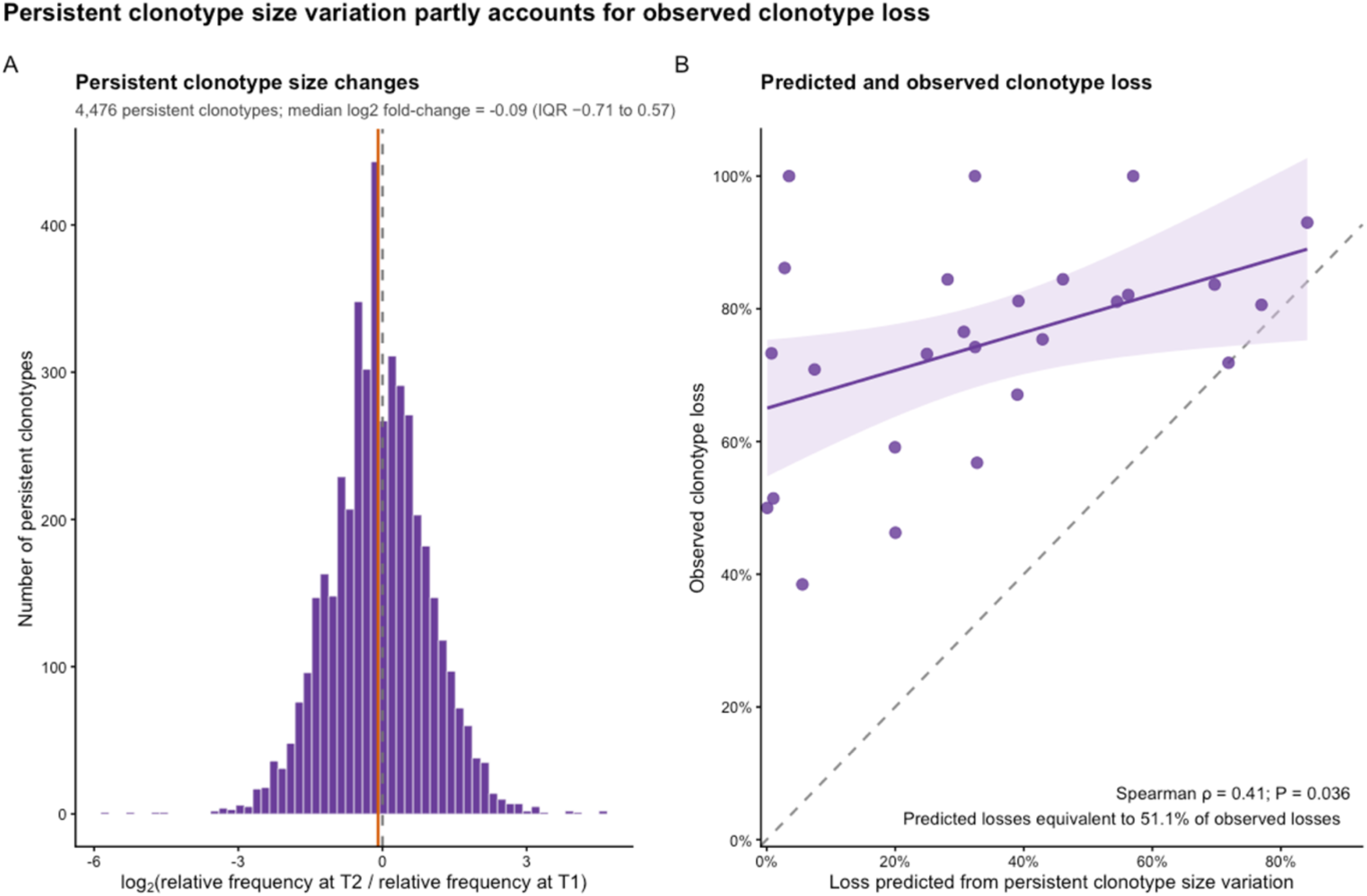
Longitudinal repertoire composition and the contribution of clonotype size variation to apparent loss. **(A)** Distribution of log₂ fold changes in relative frequency between T1 and T2 among 4,476 persistent clonotypes. The grey dashed line indicates no change and the orange dashed line indicates the median log₂ fold change of −0.09 (interquartile range −0.71 to 0.57). **(B)** Participant-level relationship between observed clonotype loss and loss predicted from the distribution of size changes among persistent clonotypes. Each point represents one participant; the solid line and shaded region show the fitted relationship and 95% confidence interval, and the dashed diagonal indicates equality between predicted and observed loss. Predicted threshold-crossing losses were equivalent to 51.1% of observed losses overall and correlated moderately with observed loss (Spearman’s ρ = 0.41, *P* = 0.036).

